# MPAthic: Quantitative Modeling of Sequence-Function Relationships for massively parallel assays

**DOI:** 10.1101/054676

**Authors:** William T. Ireland, Justin B. Kinney

## Abstract

Massively parallel assays (MPAs) are being rapidly adopted for studying a wide range of DNA, RNA, and protein sequence-function relationships. However, the software available for quantitatively modeling these relationships is severely limited. Here we describe MPAthic, a software package that enables the rapid inference of such models from a variety of MPA datasets. Using both simulated and previously published data, we show that the modeling capabilities of MPAthic greatly improve on those of existing software. In particular, only MPAthic can accurately quantify the strength of epistatic interactions. These capabilities address a major need in the analysis of MPA data.

## Background

Understanding how sequence governs function is one of the central challenges in modern biology. The classic success story in this endeavor is the cracking of the genetic code, which maps each 3-nucleotide mRNA codon to a corresponding amino acid [1]. But unlike the genetic code, which can be represented in a simple tabular form, most sequence-function relationships require a quantitative description. For instance, mutating one nucleotide in a transcriptional enhancer or one amino acid in an enzyme will typically have a quantitative, not qualitative, effect on biological function. Deciphering such quantitative sequence-function relationships has proved to be far more difficult than early success on the genetic code might have suggested.

Fortunately, a variety of massively parallel assays (MPAs), which have been developed in recent years, are providing new hope for deciphering quantitative sequence-function relationships. MPAs couple functional measurements to ultra-high-throughput DNA sequencing in ways that allow thousands to millions of sequences to be assayed in a single experiment; see [2, 3, 4, 5, 6] for reviews of these methods. Fig. 1 provides an illustration of three prototypical MPA assays: the Sort-Seq assay of Kinney et al., (2010) [7], the massively parallel reporter assay (MPRA) of Melnikov et al. [8], and the deep mutational scanning (DMS) assay of Fowler et al., (2010) [9].

**Figure 1.**
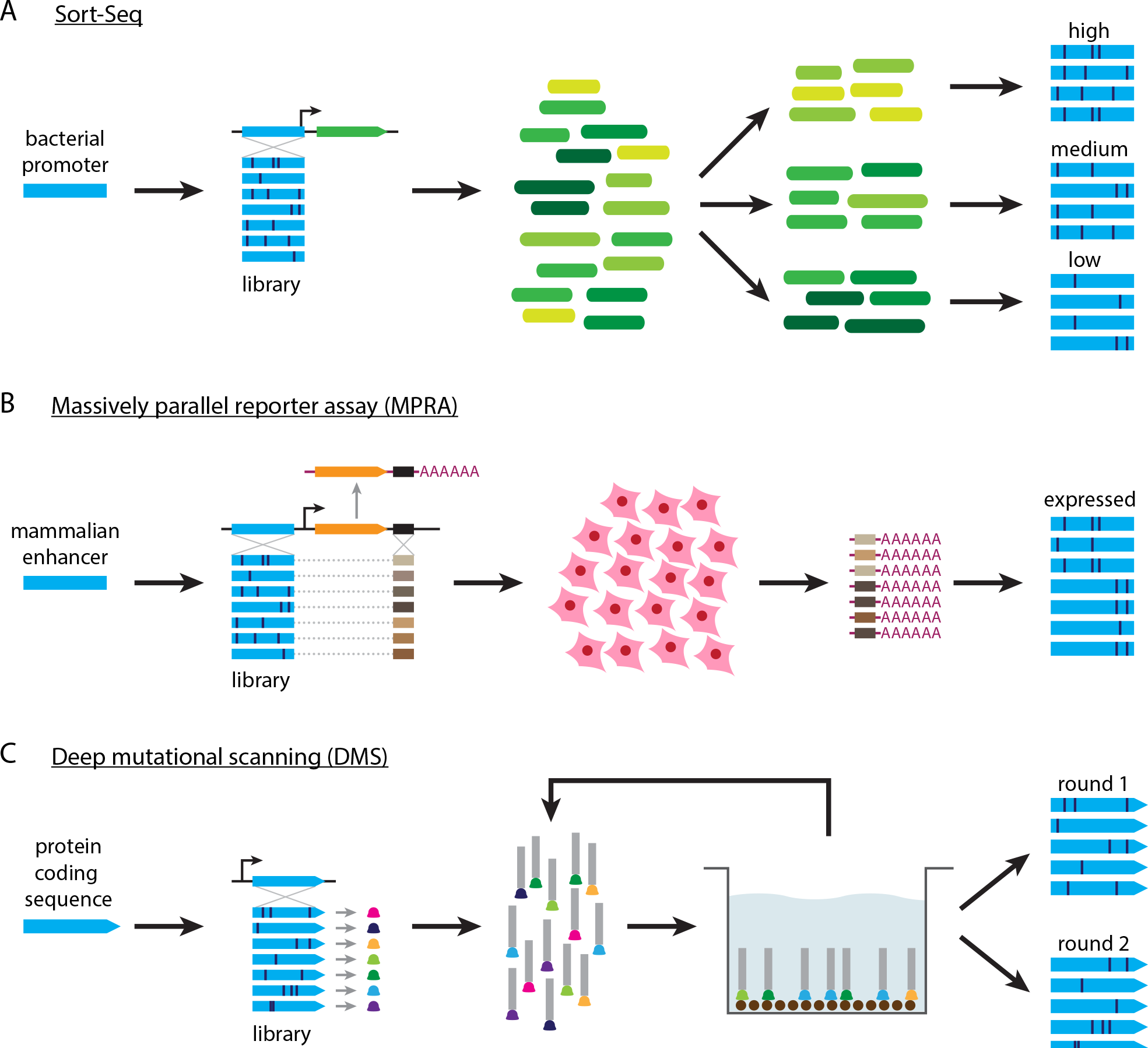
Three different massively parallel experiments. (A) The Sort-Seq assay of [7]. A plasmid library is generated in which mutagenized versions of a bacterial promoter (blue) drive the expression of a fluorescent protein (green). Cells carrying these plasmids are then sorted according to measured fluorescence using fluorescence-activated cell sorting (FACS). The variant promoters in each bin of sorted cells are then sequenced. (B) The MPRA assay of [8]. Variant enhancers (blue) are used to drive the transcription of RNA that contains enhancer-specific tags (shades of brown). Expression constructs are transfected into cell culture, after which tag-containing RNA is isolated and sequenced. Output sequences consist of the variant enhancers that correspond to expressed tags. (C) The DMS assay of [9]. Randomly mutagenized gene sequences (blue) produce variant proteins (colored bells) that are expressed on the surface of phage (gray rectangles). Panning is used to enrich for phage that express proteins that bind a specific ligand of interest (brown circles). The variant coding regions enriched after one or more rounds of panning are then sequenced.

Since the publication of these and other early works, the use of MPAs has rapidly expanded in multiple areas of molecular biology and genetics. Already, MPAs are revolutionizing the study of transcriptional regulation in vitro [10], in model organisms [7, 11, 12, 13], and in human cells [8, 14, 15], the study of splicing [16] and untranslated regions [17, 18, 19] of mRNA, the study of protein sequence-function relationships [9, 20, 21, 22], and the study of fitness landscapes and their evolutionary consequences [23, 24, 25, 26, 27]. The recent demonstration [28] of MPAs on sequence libraries integrated into mammalian chromosomes using CRISPR-Cas9 is likely to substantially increase the utility and prominence of such experiments in the near future.

Often, MPAs are used to simply catalog the activities of a large number of sequences. Multiple software packages have been developed to assist in the type of analysis that such cataloging requires [29, 30, 31]. But just tallying measurements is insufficient for characterizing most sequence-function relationships. This is because sequence space is often far too vast for every possible sequence of interest to be assayed. For example, the *σ*^70^ RNA polymerase holoenzyme of *Escherichia coli* (RNAP) has a binding site roughly 40 bp in length. The sequence specificity of RNAP is of great biological interest, and in fact RNAP was the first DNA-binding protein to have its specificity characterized [32]. But it would be impossible to store quantitative measurements for all 4^40^ ≈ 10^24^ possible binding sites, let alone make these measurements, because the resulting dataset would overwhelm the world’s information technology infrastructure [33].

Understanding a quantitative sequence-function relationship ultimately requires quantitative modeling: the construction of a mathematical function that can predict the activity of any sequence of interest regardless of whether that sequence was previously assayed in an experiment. The data that many MPAs provide is well-suited for training such models. For example, [7] used quantitative modeling of Sort-Seq data to measure the *in vivo* interaction energies governing transcriptional activation at the *lac* promoter of *E. coli*. In later work, [8] used quantitative modeling of MPRA data to design enhancers with increased inducibility in human cells.

Unfortunately, the quantitative modeling of MPA data has been severely hindered by the lack of appropriate software. The only published software that provides any quantitative modeling capabilities appropriate to MPA data is dms_tools [31]. dms_tools, however, has multiple limitations. First, it only supports the inference of ‘matrix models,’ in which each position in a sequence is assumed to contribute independently to activity.^1^ These models are blind to possible epistatic interactions between different positions within a sequence, interactions that are of great interest in MPA experiments [25]. Second, the parameters of the models that dms_tools returns are inferred using enrichment ratio calculations. Although this inference approach is common in bioinformatics, the mathematical justification for this procedure relies on multiple assumptions that are often violated in real MPA experiments [34]. Moreover, enrichment ratios can be computed only when one’s data consists of precisely one ‘selected’ set of sequences and one ‘unselected’ set. Many MPAs, e.g. [7, 9, 11, 18, 19, 22, 23, 26], yield three or more sets of sequences and it is unclear how dms_tools can be applied to these data without throwing away valuable measurements.

Here we introduce MPAthic, a software package that enables quantitative modeling of sequence-function relationships using data from a variety of MPAs. MPAthic improves on dms_tools in multiple ways. In addition to supporting the inference of matrix models using enrichment ratios, MPAthic offers two alternative inference procedures: least-squares optimization and mutual information maximization. Both of these inference methods make use of all the data produced by MPAs. Least-squares optimization facilitates the rapid analysis of MPA data, while mutual information maximization yields models that are more rigorous in theory [34] and, as will be shown, more accurate in practice. Unlike dms_tools, MPAthic also supports the inference of ‘neighbor’ models, which describe epistatic interactions between neighboring positions in a sequence.

We demonstrate MPAthic on both simulated data and on data from the MPA studies illustrated in Fig. 1 [7, 8, 9]. The modeling capabilities of MPAthic are shown to dramatically outperform those of dms_tools in almost all cases. In particular, we find that matrix models fit using mutual information maximization consistently work better than models inferred using enrichment ratio calculations. This finding validates previous theoretical arguments [34]. Moreover, we show that MPAthic can successfully infer epistatic interactions present in both simulated and real sequence-function relationships.

MPAthic incorporates other capabilities in addition to quantitative modeling: it provides methods for simulating massively parallel data using a user-specified model, for computing useful summary statistics such as ‘information footprints’ [7], and for evaluating quantitative models on arbitrary input sequences. These additional features are described in Supplemental Information (SI). All of MPAthic’s functionality is accessible at the command line. Information on obtaining MPAthic is provided in the “Availability of data and materials” section at the end of this paper.

## Methods

### Constraints on data

Many massively parallel experiments, including those of [7, 8, 9, 10, 12, 13, 17, 18, 19, 20, 21, 22, 23, 24, 25, 26, 27, 28], share the common form illustrated in Fig. 2A. One begins with a specific “wild type” sequence of interest. A library comprising variants of this wild type sequence, variants that have scattered substitution mutations, is then generated. These library sequences are then used as input to an experimental procedure that measures a specific sequence-dependent activity and, as a result of this measurement, outputs sequences into one or more “bins.” Finally, the number of occurrences of each variant sequence in each bin is assayed using high-throughput sequencing.

The resulting data consists of a set aligned sequences, each associated with a specific number of counts within each of the experimental bins. MPAthic is designed specifically for the analysis of data having this form. The aligned nature of these sequences greatly simplifies the process of quantitative modeling.

**Figure 2.**
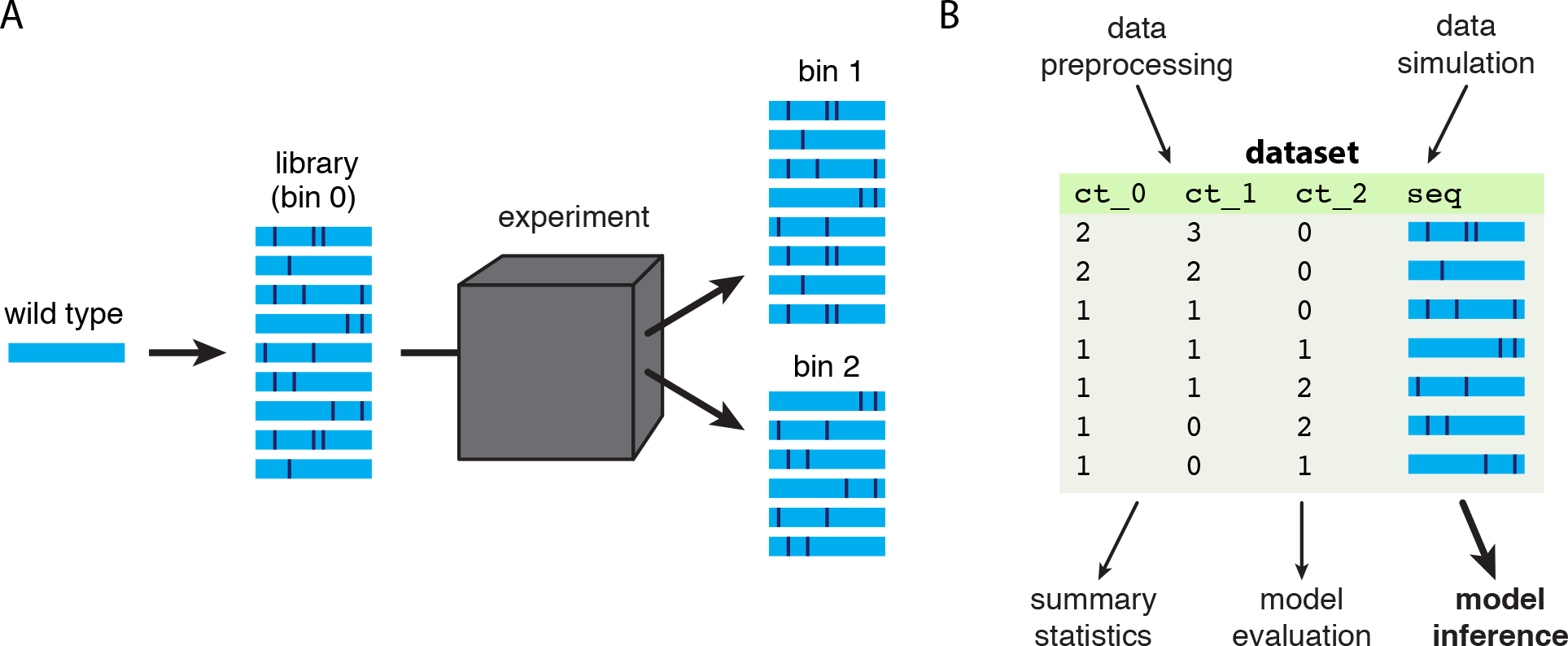
Overview of MPAthic. (A) In all massively parallel assays, a library of sequences is used as input to an experiment (black box) that outputs these sequences into one or more bins. The prevalence of each sequence in each bin depends on the assayed activity of that sequence. MPAthic can be used to analyze data from such experiments when the input library consists of substitution-mutated versions of a specific “wild type” sequence. (B) The data from such experiments can be represented as a table listing the number of occurrences of each unique sequence in each bin. MPAthic provides routines for inferring quantitative models from datasets that have this form. Routines are also provided for simulating data, for computing summary statistics, and for evaluating inferred models on arbitrary sequences.

In particular, when the assayed sequences have a natural alignment, one can sensibly model the quantitative effect of each nucleotide or amino acid position.

We note that not all massively parallel experiments have this form. SELEX-SEQ and related experiments often use library DNA that is completely random (e.g., [35, 36, 37, 38, 39, 40]), and thus cannot be expected to have well-aligned features. Other experiments, including Sort-Seq and MPRA experiments, have used libraries that contain specified arrangements of binding sites or large numbers of different genomic regions (e.g., [11, 41, 42, 43]). These other datasets are, in principle, amenable to quantitative modeling as well. This modeling task is more complex than for aligned sequences, however, and is not supported in the current version of MPAthic.

### Formalization

We formalize the problem of inferring quantitative models of sequence-function relationships as follows. We represent massively parallel data as a set of *N* sequence-measurement pairs, 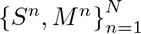 where each measurement *M^n^* is a non-negative integer corresponding to the bin in which the n’th sequence sequence, *S^n^*, was found. We assume that all sequences *S* have the same length *L*, and that at each of the *L* positions in each sequence there is one of *C* possible characters (*C* = 4 for DNA and RNA; *C* = 20 for protein). In what follows, each sequence S is represented as a binary *C* × *L* matrix having elements

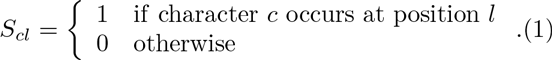

Here, *l* = 1, 2, …, *L* indexes positions within the sequence, while *c* = 1, 2, …, *C* indexes possible nucleotides or amino acids. Note that, in this representation, the same sequence *S* will typically be observed multiple times in each data set and will often fall into multiple different bins.

Our goal is to derive a function that can, given a sequence *S*, predict the activity *R* measured by the experiment. To do this we assume that the activity value *R* is given by a function *r*(*S*, *θ*) that depends on the sequence *S* and a set of parameters *θ*. Before we can infer the values of the parameters *θ* from data, we must first answer two distinct questions:

1. What functional form do we choose for *r*(*S*, *θ*)?
2. How do we use the data {*S^n^*, *M^n^*} and the model predictions *r*(*S*, *θ*) to infer parameter values *θ*?

MPAthic provides two different options for the function r: a “matrix” model, where each position in *S* contributes independently to the predicted activity, and a “neighbor” model, which accounts for potential epistatic interactions between neighboring positions. MPAthic also provides three different methods for fitting the parameters *θ* to data: parameters values can be inferred by computing enrichment ratios (a method applicable only to matrix models), by performing least squares optimization, or by maximizing the mutual information between model predictions and measurements. These different model types and inference methods are elaborated below.

### Matrix models and neighbor models

Matrix models have the form

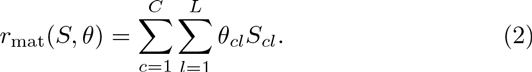

In this context, *θ* is a *C* × *L* matrix where each element *θ_cl_* represents the contribution of character *c* at position *l* to the overall sequence-dependent activity. For example, Fig. 3B shows the parameters of a matrix model that describes the sequence specificity of RNAP. These parameters were inferred from the Sort-Seq data of [7] using MPAthic.

**Figure 3.**
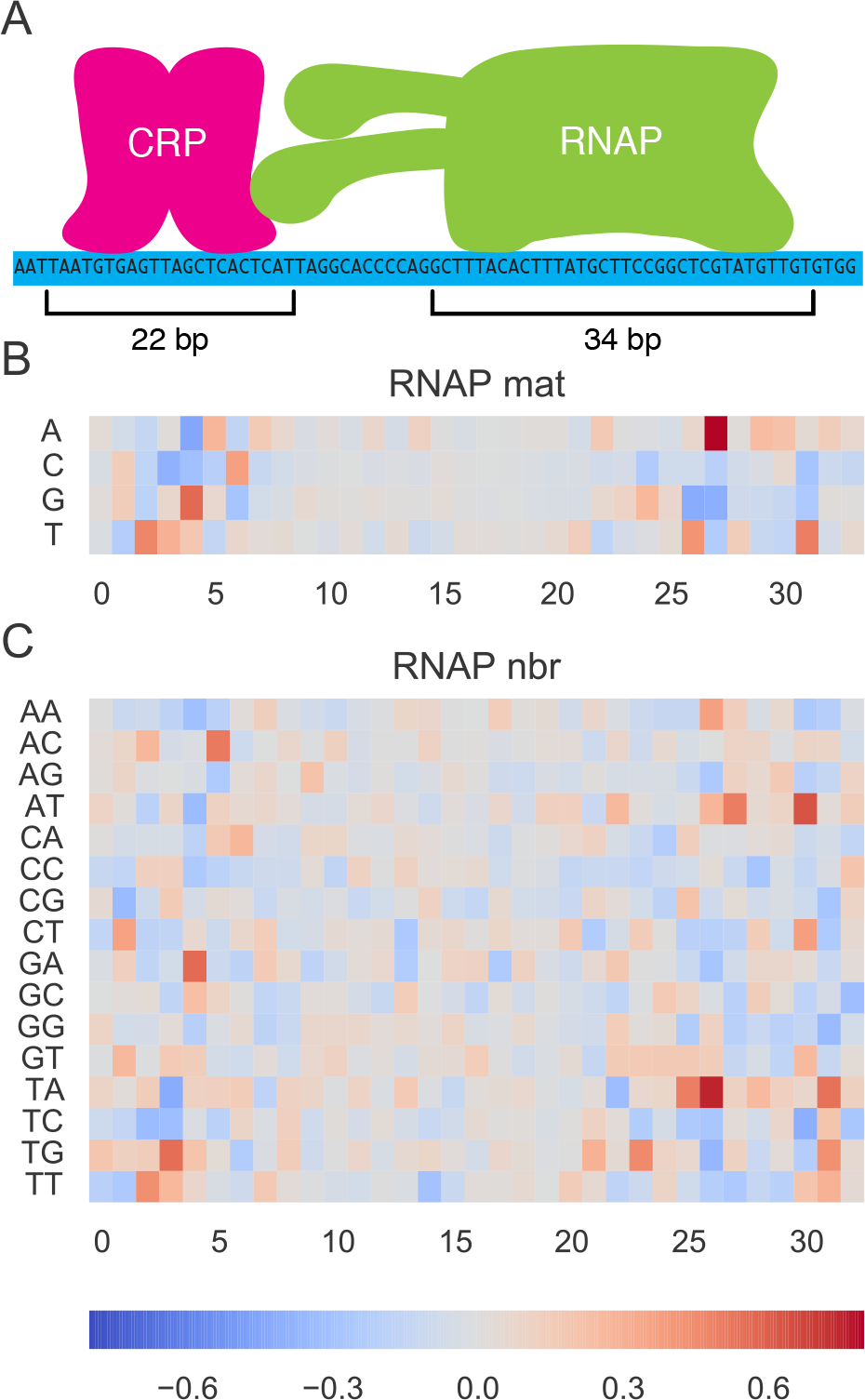
(A) The 75 bp region of the *E. coli* lac promoter that was assayed in the Sort-Seq experiments of [7]. This region contains binding sites for two proteins: CRP and RNAP. As shown in Fig. 4, multiple types of quantitative models for both CRP and RNAP (spanning the two indicated regions) were inferred from the datasets of [7] using multiple different inference methods. (B) A matrix model for RNAP, inferred from the full-wt experiment of [7] via information maximization. (C) A neighbor model for RNAP spanning the same region and fit to the same data as in panel B, again inferred using information maximization.

Neighbor models have the form

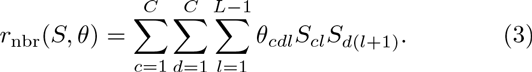

**Figure 4.**
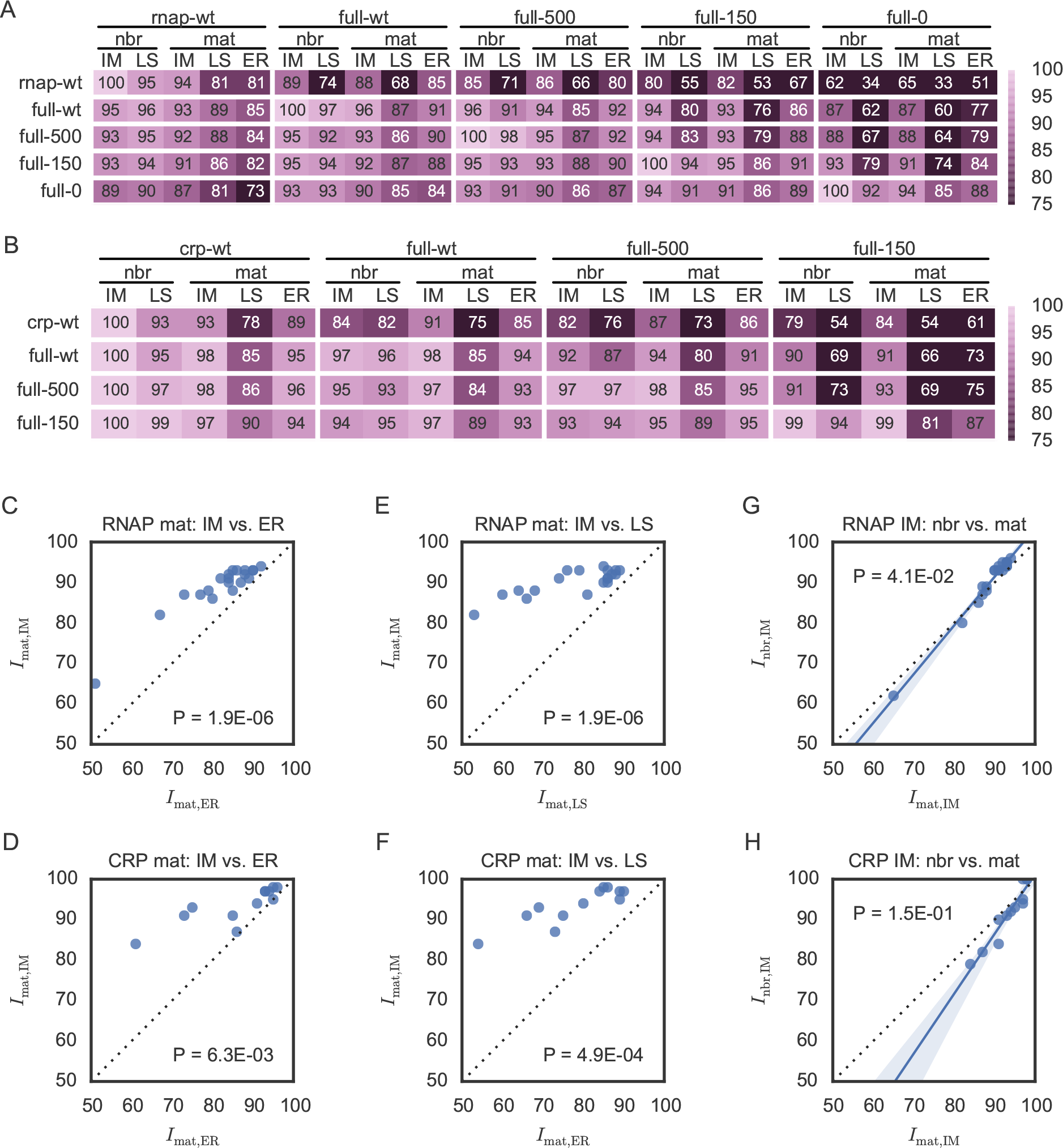
Analysis of Sort-Seq data. (A,B) Performance of (A) RNAP and (B) CRP models inferred from and evaluated on Sort-Seq data from [7]. Each column corresponds to an inferred model; column headers indicate the dataset (rnap-wt, crp-wt, full-wt, full-500, full-150, or full-0) used to train the model, the type of model inferred (neighbor (nbr) or matrix (mat)), and the inference method used by MPAthic (information maximization (IM) or least squares (LS)). Columns corresponding to matrix models inferred using dms_tools are indicated by DT. Rows indicate the datasets used to evaluate model performance. Heatmap values give the predictive information of each inferred model (column) on each test set (row). These values are expressed as a percentage of the maximal predictive information achieved on each test set (i.e., along each row). (C-H) Scatter plot comparisons of predictive information values for (C,D) matrix models fit using IM inference (1mat,IM) vs. using dms_tools (*1_mat_*,DT), (E,F) matrix models fit using IM vs. LS inference (*1_mat_*,Ls), and (G,H) IM-inferred matrix models versus IM-inferred neighbor models (*1_nbr_*,IM). Data points in panels C-H indicate model performance on non-training data only. In panels G and H, regression lines and 95% bootstrap confidence intervals are shown.

Such models comprise *C*^2^(*L* − 1) parameters, denoted *θ_cdl_*, that represent the contributions of all possible adjacent di-nucleotides or di-amino-acids within *S*. Fig. 3C illustrates one such model which, as above, describes RNAP and was inferred using MPAthic operating on data from [7].

### Enrichment ratio inference

The computation of enrichment ratios is the simplest way to infer quantitative sequence-function relationships from massively parallel data. The motivation for this inference method traces back to the seminal work of Berg and von Hippel [44, 45], and the resulting models can be thought of as the incarnation of position weight matrices [46] in the context of massively parallel experiments. We note that the calculation of enrichment ratios is one of the primary types of analyses reported in the DMS literature [2].

Enrichment ratio inference, however, places strong restrictions on the types of models and data that one can use. Specifically, one is restricted to using matrix models only, and the data used to compute parameter values can consist of only two bins of sequences: an unselected ‘library’ bin (*M* = 0) and a selected ‘enriched’ bin (*M* = 1). Moreover, the validity of this inference procedure depends on additional assumptions that are often not satisfied by real-world experiments [34].

Still, if one is willing to impose these restrictions and make the necessary assumptions, then model parameters *θ*^ER^ are computed using

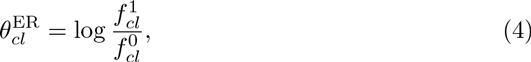

where

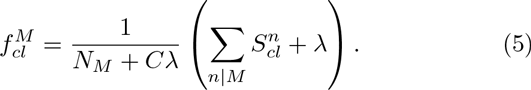

Here, 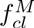 denotes the fraction of sequences in bin *M* having character *c* at position *l*, *N_M_* is the total number of sequences in bin *M*, λ is a nonnegative pseudocount (specified by the user), and the sum in Eq. (5) is restricted to sequences *S^n^* that lie within bin *M* (i.e., for which *M^n^* = *M*). We note that that Eq. (5) arises in a Bayesian calculation as a the maximum *a posteriori* estimate of 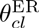 for appropriate choice of prior.

The calculation of enrichment ratios is the only type of inference supported by dms_tools [31]. This approach, however, is implemented in two different ways. dms_tools supports the computation of matrix model parameters using the formulas in Eq. (4) and Eq. (5). This leads to very rapid calculations, and for this reason such inference is also supported by MPAthic. dms_tools also supports a much more computationally intensive inference procedure, which uses Monte Carlo sampling of an explicit Bayesian posterior distribution. In [31], this Monte Carlo approach is advocated as providing a more accurate estimate of model parameters than do Eqs. (4) and (5). In what follows, we use ‘DT’ to label matrix models inferred by dms_tools using this Monte Carlo approach. Supplemental Information shows, however, that all DT models discussed in this manuscript are nearly indistinguishable from the models inferred by analytic enrichment ratio calculations.

### Least squares inference

Least squares provides a computationally simple inference procedure that overcomes the most onerous restrictions of enrichment ratio calculations. It can be used to infer any type of linear model, including both matrix models and neighbor models. It can also be used on data that consists of more than two bins.

The idea behind the least squares approach is to choose parameters *θ*^LS^ that minimize a quadratic loss function. Specifically, we use

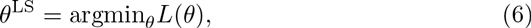

where

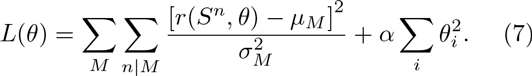

Here, *µ_M_* is the assumed mean activity of sequences in bin *M*, 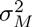 is the assumed variance in the activities of such sequences, *i* indexes all parameters in the model, and *α* is a “ridge regression” regularization parameter [47]. By using the objective function *L*(*θ*), one can rapidly compute values of the optimal parameters *θ* using standard algorithms [48].

One downside to least squares inference is the need to assume specific values for *µ_M_* and for 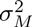 for each bin *M*. MPAthic allows the user to manually specify these values. There is a danger here, since assuming incorrect values for *µ_M_* and 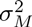 will generally lead to bias in the inferred parameters *θ*^LS^ [34]. In practice, however, the default choice of *µ_M_* = *M* and 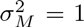 often works surprisingly well when bins are arranged from lowest to highest average activity.

Another downside to least squares is the need to assume that experimental noise - specifically, *p*(*R*|*M*) - is Gaussian. Only in such cases does least squares inference correspond to a meaningful maximum likelihood calculation. In massively parallel assays, however, noise is often strongly non-Gaussian. In such situations, least squares inference cannot be expected to yield correct model parameters for any choice of *µ_M_* and 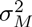 [34].

### Information maximization inference

An alternative inference procedure, one that does not suffer from the need to assume a specific quantitative form for experimental noise, is the maximization of mutual information. In the large data limit, information maximization is equivalent to performing maximum likelihood inference when the quantiative form of experimental noise is unknown [34, 49, 50]. This approach was originally proposed for receptive field inference in sensory neuroscience [51, 52, 53], but has since been applied in multiple molecular biology contexts [49, 54], including in the analysis of massively parallel experiments [7, 8].

In this approach, parameter values are chosen to maximize the mutual information between model predictions and measurements. Specifically, one chooses

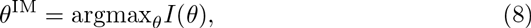

where

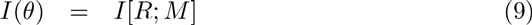

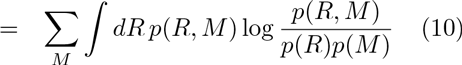

is the mutual information between the bins *M* in which sequences are found and the corresponding model prediction *R* for those sequences. In what follows, *I*(*θ*) is referred to as the “predictive information” of the model. For each choice of *θ*, computing predictive information requires a regularized estimate of the joint probability distribution *p*(*R*, *M*). MPAthic currently uses standard kernel density methods [47] to estimate these distributions, although field-theoretic density estimation [55, 56] may ultimately prove superior in this context.

Following [7], MPAthic optimizes parameters using a Metropolis Monte Carlo algorithm in which each choice for *θ* has relative probability exp[*NI*(*θ*)]. Because this Monte Carlo procedure is computationally expensive, information maximization is much slower than enrichment ratio calculations or least squares inference. Running on a standard laptop computer, our current algorithm takes between 30 minutes and 2 hours for each of the information maximization tasks described below.

## Results

To test the capabilities of MPAthic, we analyzed data from previously published Sort-Seq [7], MPRA [8], and DMS [9] studies. Each of these studies generated multiple independent datasets, allowing us to test the inference capabilities of MPAthic by training and testing models on separate data. We also analyzed simulated data in order to assess the ability of MPAthic to accurately recover known parameter values.

### Sort-Seq data

In their studies of the *E. coli lac* promoter, Kinney et al. [7] performed six independent Sort-Seq experiments, which they referred to as rnap-wt, crp-wt, full-wt, full-500, full-150, and full-0. All of these experiments assayed the transcriptional activity of variant sequences spanning a 75 bp region of the *lac* promoter (Fig. 3A). This region is known to bind two proteins, RNAP and CRP, at two separate binding sites. In the original study [7], models for the sequence specificity of these two proteins were inferred by modeling how transcription depends on sequence variation within their respective binding sites.

For both RNAP and CRP, we used each of these six datasets to infer both matrix models and neighbor models.^2^ Inference was performed using two methods supported by MPAthic: least squares optimization (LS) and mutual information maximization (IM). In addition, matrix models were inferred using the Monte Carlo estimates of enrichment ratios provided by dms_tools (DT).^3^ The performance of each of these models on each of the available datasets was then quantified by estimating the predictive information *I*[*R*; *M*].

Fig. 4A illustrates the performance of each inferred RNAP model (columns) on each of the published datasets (rows). Fig. 4B shows similar results for the inferred CRP models.^4^ For both CRP and RNAP, the IM-inferred matrix models consistently outperformed the LS- and DT-inferred matrix models when evaluated on independent test data (Figs. 4C,4D,4E,4F). This finding lends support to the theoretical arguments of [34] that information maximization has substantial advantages over other methods for inferring quantitative sequence-function relationships from massively parallel data. It also demonstrates the superior performance of MPAthic relative to dms_tools for the inference of matrix models.

We also investigated whether neighbor models, which account for epistatic interactions between neighboring positions in a sequence, might provide better descriptions of RNAP and CRP than simple matrix models do. To our knowledge, the presence of such interactions in either of these well-studied proteins has yet to be definitively established (although see [57]). We therefore compared the predictive performance of matrix and neighbor models that were trained (using IM) on the same datasets (Figs. 4G, 4H).

Neighbor models did not always outperform matrix models in these tests. However, for both CRP and RNAP, neighbor models did perform better than their corresponding matrix models when the predictive information of the matrix model was high (Figs. 4E,4F). Such high matrix model predictive information values are expected to occur when the data used to train models is of high quality. We interpret this finding as evidence for epistatic interactions in the specificities of both CRP and RNAP. This also indicates that MPAthic, but not dms_tools, is capable of quantifying such interactions. We note that the crp-wt data should be expected to be the best dataset for training models of CRP because the mutation rate used in this experiment was the highest (24%). This expectation is consistent with our finding that the CRP neighbor model inferred from crp-wt outperformed every other model inferred for CRP.

### Simulated data

To further establish the ability of MPAthic to properly infer quantitative models, we next analyzed simulated Sort-Seq data. To generate simulated Sort-Seq data, we used the simulation capabilities of MPAthic together with the nbr-IM models for RNAP and CRP that were inferred from the full-wt dataset of [7]. Eight datasets were simulated in total, four for RNAP and four for CRP. In each simulation, 10^6^ cells were sorted into either 10 or 2 bins; see SI for simulation details. Half of these simulated datasets (labeled “train”) were then used to infer matrix and neighbor models as described in the previous section. The other half (labeled “test”) were used solely to evaluate model performance.

Fig. 5A shows results for the simulated RNAP data, while Fig. 5B shows corresponding results for simulated CRP data. The nbr-IM models performed best in every case tested, with virtually no apparent difference in performance between training and test data. In particular, all of the nbr-IM models performed substantially better than the mat-IM models, demonstrating the ability of MPAthic to accurately quantify epistatic interactions. It is also worth noting that, as in the analysis of Sort-Seq data, the matrix models found by MPAthic using IM inference outperformed those computed by dms_tool using enrichment ratio calculations. Figs. 5C and 5D plot the values of parameters for inferred neighbor models against the corresponding parameter values of the neighbor models used to generate the data. We found very strong agreement, with a signal-to-noise ratio of 31 across the 528 parameters of the RNAP neighbor model, and a signal-to-noise ratio of 49 across the 336 parameters of the CRP neighbor model.

**Figure 5.**
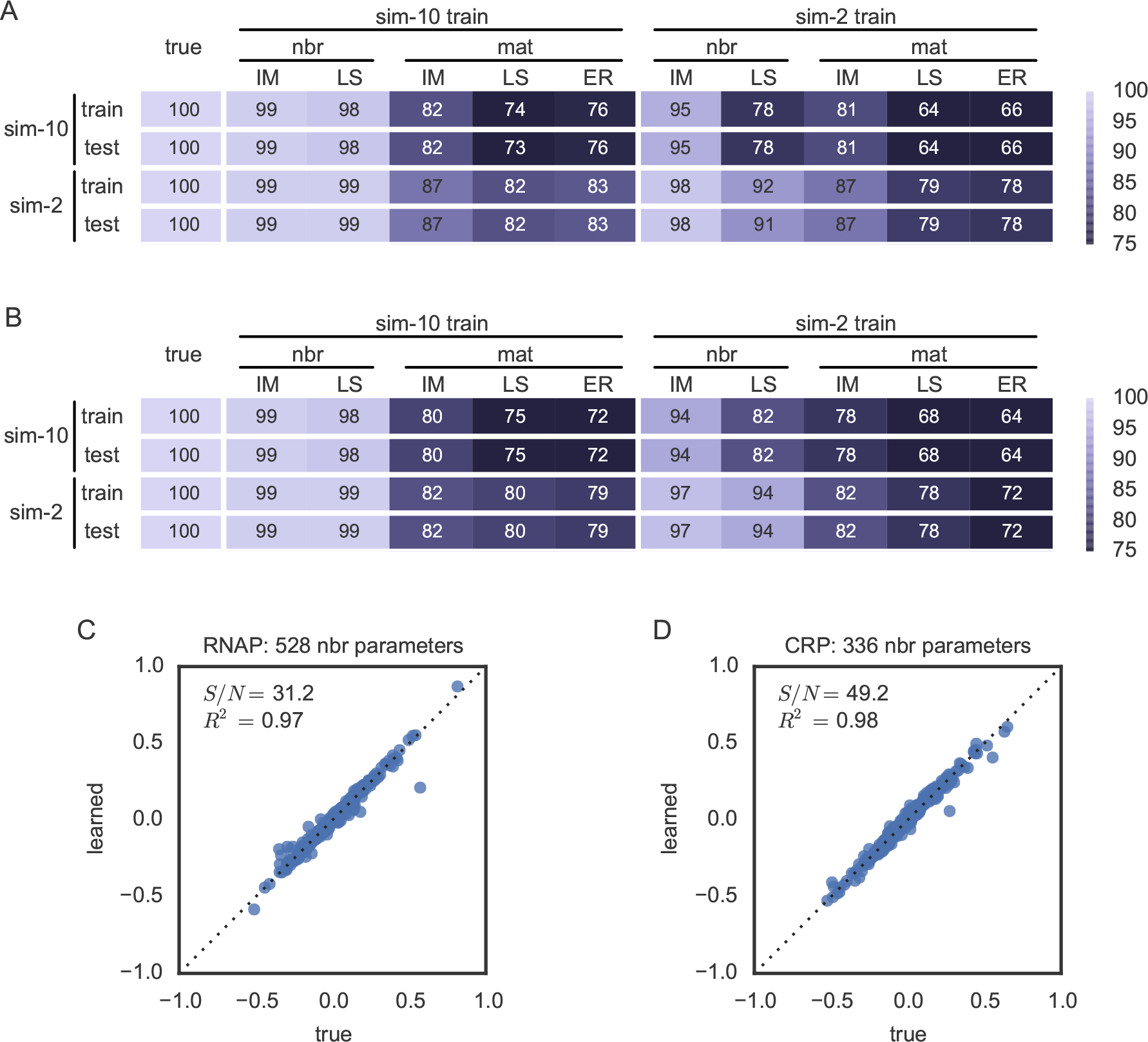
Analysis of simulated data. Sort-Seq data was simulated using the RNAP and CRP neighbor models inferred using MPAthic (in IM mode) from the full-wt data of [7]. Four datasets were generated for each model: one training and one test set were generated by sorting into 10 bins, while one training and one test set generated by sorting into 2 bins. (A,B) Performance of (A) RNAP and (B) CRP models inferred from and evaluated on these simulated datasets. Columns indicate the dataset used to train the model, the type of model inferred (nbr or mat), and the inference method used for training (MPAthic in IM or LS mode, or dms_tools (DT)). Rows indicate the datasets on which models were evaluated. As in Figs. 3A and 3B, heatmaps show predictive information values expressed as a percentage of the maximal predictive information achieved on each dataset. (C,D) Comparison of the parameters of the neighbor models used in these simulations to the parameters of the neighbor models fit to the corresponding “sim-10 train” data using MPAthic in IM mode. Also shown is the signal-to-noise ratio, defined as the variance in the abscissa divided by the variance in the deviation of the ordinate from the diagonal.

### MPRA and DMS data

MPAthic is designed to facilitate the quantitative modeling of data from a variety of massively parallel assays, including MPRA and DMS experiments. To test the utility of MPAthic in these contexts, we inferred matrix models using MPRA data from [8] and DMS data from [9].^5^

In [8], replicate MPRA experiments were performed on a synthetic cAMP responsive element (CRE). These experiments tested ~ 2.7 × 10^4^ microarray-synthesized CREs having randomly scattered substitution mutations (10% per nucleotide position) throughout an 87 bp region. Using MPAthic, we inferred matrix models spanning this entire 87 bp region using IM- and LS-based inference. We also computed matrix models using dms_tools. We found that on both replicate datasets, the IM-inferred models found by MPAthic performed the best in cross-comparisons (Fig. 6A). Moreover, both of these models performed better on both replicate datasets relative to the matrix model described in the original publication [8].

**Figure 6.**
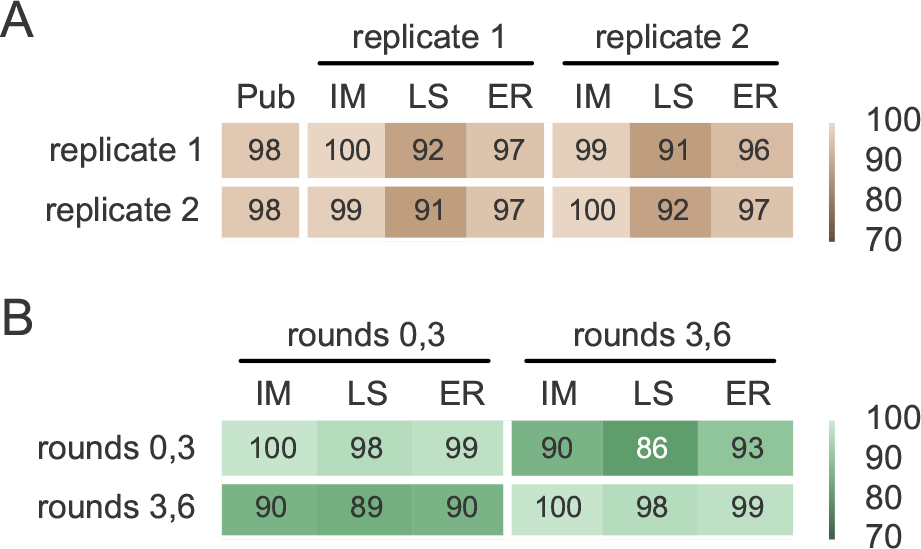
Analysis of MPRA and DMS data. (A)Cross-comparison of matrix models fit to data from two replicate MPRA experiments reported in [8]. The performance of the matrix model reported in the original publication (Pub) is also shown. (B) In the DMS experiments of [9], sequence data was gathered after 0, 3, and 6 rounds of selection. Shown is a cross-comparison of matrix models fit to data from either rounds 0 and 3, or to data from rounds 3 and 6, using either MPAthic in IM or LS mode, or using dms_tools (DT).x

The DMS experiments of [9] assayed a variable region spanning 33 aa within a WW domain protein. Specifically, the gene sequences of this WW domain was mutagenized at ~2% per base. Multiple rounds of panning using a peptide ligand were then used to select WW-domain variants displayed on the surface of phage. The WW domain coding sequences present in the phage library after 0, 3, and 6 rounds of selection were then sequenced.

Using MPAthic, we fit models to either the round 0 and round 3 libraries, or to the round 3 and round 6 libraries. When trained on round 0,3 data and tested on round 3,6 data, the IM-inferred matrix models returned by MPAthic performed better than LS-inferred models and about the same as the matrix models returned by dms_tools (Fig. 6B). However, IM models fit to round 3,6 data actually performed worse than the corresponding models of dms_tools. This is the only situation we encountered where DT models outperformed IM models.

The poor performance of IM in this context is most likely due to the sparsity of data in the round 3,6 dataset. Specifically, in the round 3,6 dataset, we observed 8 amino-acid-position combinations with no representation in the data. Furthermore, 16 amino-acid-position combinations were represented by data from only one sequence read. By contrast, the round 0,3 dataset contained data on all amino-acid-position combinations, and for only 2 of these combinations did this data come from a single sequence. Our results therefore suggest that the IM-based inference of MPAthic can perform at least as well on DMS data as enrichment ratio calculations, but only when datasets are sufficiently rich. More generally, the existence of 20 amino acids compared to 4 DNA/RNA bases places a significantly larger burden on the amount of data needed to obtain accurate models from DMS data relative to Sort-Seq or MPRA data. This is true regardless of the inference method one uses.

## Discussion

MPAthic provides routines for inferring quantitative models from MPA data. Such modeling is essential for understanding quantitative sequence-function relationships. The lack of published software available for this purpose has likely restricted the range of biological problems to which MPAs have been applied. Currently, the only published software for learning quantitative models from MPA data is dms_tools [31]. As shown here, MPAthic improves upon dms_tools in two key ways.

First, MPAthic is better than dms_tools at inferring the parameters of matrix models, the simplest and most widely used type of model for describing sequence-function relationships. This improved performance is due to MPAthic supporting the use of mutual information maximization as a way to infer parameter values. dms_tools, by contrast, is limited to enrichment ratio calculations. Mutual information maximization has been theoretically shown to provide an optimal inference method in the large data limit [34]. By contrast, the use of enrichment ratio calculations requires multiple assumptions that are often violated in real-world MPA experiments. Moreover, mutual information maximization makes use of all the available data, while enrichment ratio calculations often require one to discard valuable measurements. As we showed on both real and simulated data, the mutual information maximization routines provided by MPAthic almost always yield better matrix models than do enrichment ratio calculations. The only exception to this observation was found, unsurprisingly, in the analysis of a dataset having comparatively sparse coverage.

Second, unlike dms_tools, MPAthic enables the quantification of epistatic interactions via the inference of neighbor models. Using simulated data, we showed that MPAthic is able to recover nearest-neighbor epistatic interactions with high accuracy. When applied to the published Sort-Seq data of [7], MPAthic was able to find neighbor models for both RNAP and CRP that had higher predictive power than the corresponding optimal matrix models. This indicates the successful quantification of real epistatic interactions that had not been previously known for either of these well-studied proteins.

The quantitative modeling of sequence-function relationships will ultimately require capabilities beyond those currently supported by MPAthic. For instance, there are a variety of other types of models that are likely to prove useful. Particularly promising are models with sparse all-versus-all pairwise interactions [57], models with interactions based on higher-order sequence features [58], deep neural network models [59], and nonlinear models that reflect specific biophysical mechanisms [7]. MPAthic does, however, provide a framework into which such modeling capabilities can be incorporated in the future, and through which different modeling strategies can be compared in a transparent way.

## Availability of data and materials

- Project: MPAthic (v0.01.01)
- Homepage: github.com/jbkinney/mpathic
- Archived version: DOI 10.5281/zenodo.55837
- Operating systems: Platform independent
- Programming language: Python, Cython
- Installation

– Clone the GitHub repository and install using ‘python setup.py install’.
– Install via PyPI using ‘pip install mpathic’.
- Other requirements: POSIX command line, Python 2.7.9, multiple Python packages.
- License: New BSD
- Restrictions on use by non-academics: None.

## Author’s contributions

WTI and JBK participated in all aspects of this research.

## Footnotes

1 We use the term ‘matrix model’ here instead of the more common term ‘position weight matrix’ to distinguish the mathematical form of such models from the method by which model parameters are inferred from data. The term ‘position-specific scoring matrix’ (PSSM) is synonymous with ‘matrix model.’

2 Raw data from [7] is available on NCBI SRA, accession number SRA012345; processed data formatted foruse with MPAthic is provided on the MPAthic project homepage.

3 Because enrichment ratio calculations require exactly two sequence bins, only one high-activity and one low-activity set of sequences were used for the enrichment ratio calculations in Figs. 4 and 5.

4 RNAP models were not trained or tested on the crp-wt dataset because the RNAP binding site was not mutagenized in that experiment. Similarly, CRP models were not trained or tested on the rnap-wt dataset. CRP models were also not trained or tested on the full-0 dataset because cAMP, a ligand that CRP requires in order to bind DNA, was absent in that experiment.

5 The preprocessed MPRA data of [8] was obtained from NCBI GEO, accession number GSE31982. The preprocessed DMS data of [2] was kindly provided by Douglas Fowler; raw data is available from NCBI SRA, accession number SRA020603. Processed data frommboth publications, formatted for use with MPAthic, is provided on the MPAthic project homepage. The neighbor models fit to data from both of these studies performed poorly relative to matrix models. We therefore ignore these neighbor models in what follows.

## Acknowledgements

We thank Douglas Fowler for providing preprocessed DMS data from [9]. We also thank Tarjei Mikkelsen for posting the preprocessed MPRA data from [8] on NCBI GEO. WTI was supported by NIH grants DP1 OD000217 and R01 GM085286. The work of JBK was supported by the Simons Center for Quantitative Biology at Cold Spring Harbor Laboratory.

## References

1. Judson, H.F.: The Eighth Day of Creation. Cold Spring Harbor Laboratory, Cold Spring Harbor (1996)

2. Fowler, D.M., Fields, S.: Deep mutational scanning: a new style of protein science. Nat Methods 11(8), 801–807 (2014)

3. White, M.A.: Understanding how cis-regulatory function is encoded in DNA sequence using massively parallel reporter assays and designed sequences. Genomics 106(3), 165–170 (2015)

4. Inoue, F., Ahituv, N.: Decoding enhancers using massively parallel reporter assays. Genomics 106(3), 159–164 (2015)

5. Shendure, J., Fields, S.: Massively Parallel Genetics. Genetics 203(2), 617–619 (2016)

6. Peterman, N., Levine, E.: Sort-seq under the hood: implications of design choices on large-scale characterization of sequence-function relations. BMC Genomics 17(1), 206 (2016)

7. Kinney, J.B., Murugan, A., Callan, C.G., Cox, E.C.: Using deep sequencing to characterize the biophysical mechanism of a transcriptional regulatory sequence. Proc Natl Acad Sci USA 107(20), 9158–9163 (2010)

8. Melnikov, A., Murugan, A., Zhang, X., Tesileanu, T., Wang, L., Rogov, P., Feizi, S., Gnirke, A., Callan, C.G., Kinney, J.B., Kellis, M., Lander, E.S., Mikkelsen, T.S.: Systematic dissection and optimization of inducible enhancers in human cells using a massively parallel reporter assay. Nat Biotechnol 30(3), 271–277 (2012)

9. Fowler, D.M., Araya, C.L., Fleishman, S.J., Kellogg, E.H., Stephany, J.J., Baker, D., Fields, S.: High-resolution mapping of protein sequence-function relationships. Nat Methods 7(9), 741–746 (2010)

10. Patwardhan, R.P., Lee, C., Litvin, O., Young, D.L., Pe'er, D., Shendure, J.: High-resolution analysis of DNA regulatory elements by synthetic saturation mutagenesis. Nat Biotechnol 27(12), 1173–1175 (2009)

11. Sharon, E., Kalma, Y., Sharp, A., Raveh-Sadka, T., Levo, M., Zeevi, D., Keren, L., Yakhini, Z., Weinberger, A., Segal, E.: Inferring gene regulatory logic from high-throughput measurements of thousands of systematically designed promoters. Nat Biotechnol 30(6), 521–530 (2012)

12. Patwardhan, R.P., Hiatt, J.B., Witten, D.M., Kim, M.J., Smith, R.P., May, D., Lee, C., Andrie, J.M., Lee, S.-I., Cooper, G.M., Ahituv, N., Pennacchio, L.A., Shendure, J.: Massively parallel functional dissection of mammalian enhancers in vivo. Nat Biotechnol 30(3), 265–270 (2012)

13. Kwasnieski, J.C., Mogno, I., Myers, C.A., Corbo, J.C., Cohen, B.A.: Complex effects of nucleotide variants in a mammalian cis-regulatory element. Proc Natl Acad Sci USA 109(47), 19498–19503 (2012)

14. Ulirsch, J.C., Nandakumar, S.K., Wang, L., Giani, F.C., Zhang, X., Rogov, P., Melnikov, A., McDonel, P., Do, R., Mikkelsen, T.S., Sankaran, V.G.: Systematic Functional Dissection of Common Genetic Variation Affecting Red Blood Cell Traits. Cell 165(6), 1530–1545 (2016)

15. Tewhey, R., Kotliar, D., Park, D.S., Liu, B., Winnicki, S., Reilly, S.K., Andersen, K.G., Mikkelsen, T.S., Lander, E.S., Schaffner, S.F., Sabeti, P.C.: Direct Identification of Hundreds of Expression-Modulating Variants using a Multiplexed Reporter Assay. Cell 165(6), 1519–1529 (2016)

16. Rosenberg, A.B., Patwardhan, R.P., Shendure, J., Seelig, G.: Learning the Sequence Determinants of Alternative Splicing from Millions of Random Sequences. Cell 163(3), 698–711 (2015)

17. Holmqvist, E., Reimegård, J., Wagner, E.G.H.: Massive functional mapping of a 5'-UTR by saturation mutagenesis, phenotypic sorting and deep sequencing. Nucl Acids Res 41(12), 122 (2013)

18. Peterman, N., Lavi-Itzkovitz, A., Levine, E.: Large-scale mapping of sequence-function relations in small regulatory RNAs reveals plasticity and modularity. Nucl Acids Res 42(19), 12177–12188 (2014)

19. Oikonomou, P., Goodarzi, H., Tavazoie, S.: Systematic identification of regulatory elements in conserved 3’ UTRs of human transcripts. Cell Rep 7(1), 281–292 (2014)

20. Whitehead, T.A., Chevalier, A., Song, Y., Dreyfus, C., Fleishman, S.J., De Mattos, C., Myers, C.A., Kamisetty, H., Blair, P., Wilson, I.A., Baker, D.:Optimization of affinity, specificity and function of designed influenza inhibitors using deep sequencing. Nat Biotechnol30(6), 543–548 (2012)

21. McLaughlin, R.N., Poelwijk, F.J., Raman, A., Gosal, W.S., Ranganathan, R.: The spatial architecture of protein function and adaptation. Nature 491(7422), 138–142 (2012)

22. Adams, R.M., Kinney, J.B., Mora, T., Walczak, A.M.: Measuring the sequence-affinity landscape of antibodies with massively parallel titration curves. bioRxiv (2016). related:ejt9xXT9yLYJ

23. Hietpas, R.T., Jensen, J.D., Bolon, D.N.A.: Experimental illumination of a fitness landscape. Proc Natl Acad Sci USA 108(19), 7896–7901 (2011)

24. Thyagarajan, B., Bloom, J.D.: The inherent mutational tolerance and antigenic evolvability of influenza hemagglutinin. Elife 3 (2014)

25. Podgornaia, A.I., Laub, M.T.: Pervasive degeneracy and epistasis in a protein-protein interface. Science 347(6222), 673–677 (2015)

26. Sarkisyan, K.S., Bolotin, D.A., Meer, M.V., Usmanova, D.R., Mishin, A.S., Sharonov, G.V., Ivankov, D.N., Bozhanova, N.G., Baranov, M.S., Soylemez, O., Bogatyreva, N.S., Vlasov, P.K., Egorov, E.S., Logacheva, M.D., Kondrashov, A.S., Chudakov, D.M., Putintseva, E. V., Mamedov, I.Z., Tawfik, D.S., Lukyanov, K.A., Kondrashov, F.A.: Local fitness landscape of the green fluorescent protein. Nature 533(7603), 397–401 (2016)

27. Li, C., Qian, W., Maclean, C.J., Zhang, J.: The fitness landscape of a tRNA gene. Science 352(6287), 837–840 (2016)

28. Findlay, G.M., Boyle, E.A., Hause, R.J., Klein, J.C., Shendure, J.: Saturation editing of genomic regions by multiplex homology-directed repair. Nature 513(7516), 120–123 (2014)

29. Fowler, D.M., Araya, C.L., Gerard, W., Fields, S.: Enrich: software for analysis of protein function by enrichment and depletion of variants. Bioinformatics 27(24), 3430–3431 (2011)

30. Alam, K.K., Chang, J.L., Burke, D.H.: FASTAptamer: A Bioinformatic Toolkit for High-throughput Sequence Analysis of Combinatorial Selections. Mol Ther Nucleic Acids 4(3), 230 (2015)

31. Bloom, J.D.: Software for the analysis and visualization of deep mutational scanning data. BMC Bioinformatics 16, 168 (2015)

32. Pribnow, D.: Nucleotide sequence of an RNA polymerase binding site at an early T7 promoter. Proc Natl Acad Sci USA 72(3), 784–788 (1975)

33. Hilbert, M., López, P.: The world’s technological capacity to store, communicate, and compute information. Science 332(6025), 60–65 (2011)

34. Atwal, G.S., Kinney, J.B.:Learning Quantitative Sequence-Function Relationships from Massively Parallel Experiments. J Stat Phys 162(5), 1203–1243 (2016)

35. Zykovich, A., Korf, I., Segal, D.J.: Bind-n-Seq: high-throughput analysis of in vitro protein-DNA interactions using massively parallel sequencing. Nucl Acids Res 37(22), 151–151 (2009)

36. Zhao, Y., Granas, D., Stormo, G.D.: Inferring binding energies from selected binding sites. PLoS Comput Biol 5(12), 1000590 (2009)

37. Jolma, A., Kivioja, T., Toivonen, J., Cheng, L., Wei, G., Enge, M., Taipale, M., Vaquerizas, J.M., Yan, J., Sillanpää, M.J., Bonke, M., Palin, K., Talukder, S., Hughes, T.R., Luscombe, N.M., Ukkonen, E., Taipale, J.: Multiplexed massively parallel SELEX for characterization of human transcription factor binding specificities. Genome Res 20(6), 861–873 (2010)

38. Wong, D., Teixeira, A., Oikonomopoulos, S., Humburg, P., Lone, I.N., Saliba, D., Siggers, T., Bulyk, M., Angelov, D., Dimitrov, S., Udalova, I.A., Ragoussis, J.: Extensive characterization of NF-kB binding uncovers non-canonical motifs and advances the interpretation of genetic functional traits. Genome Biol 12(7), 70 (2011)

39. Slattery, M., Riley, T., Liu, P., Abe, N., Gomez-Alcala, P., Dror, I., Zhou, T., Rohs, R., Honig, B., Bussemaker, H.J., Mann, R.S.: Cofactor binding evokes latent differences in DNA binding specificity between Hox proteins. Cell 147(6), 1270–1282 (2011)

40. Jolma, A., Yan, J., Whitington, T., Toivonen, J., Nitta, K.R., Rastas, P., Morgunova, E., Enge, M., Taipale, M., Wei, G., Palin, K., Vaquerizas, J.M., Vincentelli, R., Luscombe, N.M., Hughes, T.R., Lemaire, P., Ukkonen, E., Kivioja, T., Taipale, J.: DNA-binding specificities of human transcription factors. Cell 152(1–2), 327–339 (2013)

41. Smith, R.P., Taher, L., Patwardhan, R.P., Kim, M.J., Inoue, F., Shendure, J., Ovcharenko, I., Ahituv, N.: Massively parallel decoding of mammalian regulatory sequences supports a flexible organizational model. Nat Genet 45(9), 1021–1028 (2013)

42. Kheradpour, P., Ernst, J., Melnikov, A., Rogov, P., Wang, L., Zhang, X., Alston, J., Mikkelsen, T.S., Kellis, M.: Systematic dissection of regulatory motifs in 2000 predicted human enhancers using a massively parallel reporter assay. Genome Res 23(5), 800–811 (2013)

43. Mogno, I., Kwasnieski, J.C., Cohen, B.A.: Massively parallel synthetic promoter assays reveal the in vivo effects of binding site variants. Genome Res 23(11), 1908–1915 (2013)

44. Berg, O., von Hippel, P.: Selection of DNA binding sites by regulatory proteins. Statistical-mechanical theory and application to operators and promoters. J Mol Biol 193(4), 723–750 (1987)

45. Berg, O., von Hippel, P.: Selection of DNA binding sites by regulatoryproteins. II. The binding specificity of cyclic AMP receptor protein to recognition sites. J Mol Biol 200(4), 709–723 (1988)

46. Stormo, G., Fields, D.: Specificity, free energy and information content in protein-DNA interactions. Trends Biochem Sci 23(3), 109–113(1998)

47. Hastie, T., Tibshirani, R., Friedman, J.: The Elements of Statistical Learning, 2nd edn. Springer, New York (2011)

48. Press, W., Teukolsky, S., Wetterling, W., Flannery, B.: Numerical Recipes in C: the Art of Scientific Computing. Cambridge University Press, Cambridge (1997)

48. Kinney, J.B., Tkacik, G., Callan, C.G.: Precise physical models of protein-DNA interaction from high-throughput data. Proc Natl Acad Sci USA 104(2), 501–506 (2007)

50. Kinney, J.B., Atwal, G.S.: Parametric inference in the large data limit using maximally informative models. Neural Comput 26(4), 637–653 (2014)

51. Sharpee, T., Rust, N., Bialek, W.: Analyzing neural responses to natural signals: maximally informative dimensions. Neural Comput 16(2), 223–250 (2004)

52. Paninski, L.:Convergence properties of three spike-triggered analysis techniques. Network-Comp Neural 14(3), 437–464 (2003)

53. Sharpee, T., Sugihara, H., Kurgansky, A., Rebrik, S., Stryker, M., Miller, K.: Adaptive filtering enhances information transmission in visual cortex. Nature 439(7079), 936–942 (2006)

54. Elemento, O., Slonim, N., Tavazoie, S.: A universal framework for regulatory element discovery across all genomes and data types. Mol Cell 28(2), 337–350 (2007)

55. Kinney, J.B.: Estimation of probability densities using scale-free field theories. Phys Rev E, 011301 (2014)

56. Kinney, J.B.: Unification of field theory and maximum entropy methods for learning probability densities. Phys Rev E 92(3–1), 032107 (2015)

57. Otwinowski, J., Nemenman, I.: Genotype to phenotype mapping and the fitness landscape of the E. coli lac promoter. PLoS ONE 8(5), 61570 (2013)

58. Sharon, E., Lubliner, S., Segal, E.: A feature-based approach to modeling protein-DNA interactions. PLoS Comput Biol 4(8), 1000154 (2008)

59. Alipanahi, B., Delong, A., Weirauch, M.T., Frey, B.J.: Predicting the sequence specificities of DNA- and RNA-binding proteins by deep learning. Nat Biotechnol 33(8), 831–838 (2015)

